# Antimicrobial effect of two endemic Dominican plants on microorganisms that cause otitis

**DOI:** 10.1101/2024.04.09.588785

**Authors:** Julio A. Rivas-Ramírez, Michael M. Morales-Mercedes, Juan A. Guzmán-Recio, María I. Infante, Alexander Valdez-Disla, Anel E. Guzmán-Marte, Maritza Ramírez, Dolores Mejía-De La Cruz, Manuel Vásquez-Tineo

**Affiliations:** Natural Substances Research Laboratory, Autonomous University of Santo Domingo, Santo Domingo, RD; Research and Clinical Laboratory Management, Hospital General de la Plaza de la Salud, Santo Domingo, RD

**Keywords:** Ethnomedicine, Organic extracts, Endemic plants, Salvia, Otitis, Antimicrobial activity, Bacterial resistance, Phytochemical screening, Thin-layer chromatography, American type culture collection

## Abstract

**Introduction:** Otitis is a condition that typically results from infectious agents causing inflammation of the ear and its surrounding tissues. Common treatment involves the use of antimicrobial drugs to combat the infection and alleviate symptoms. However, the overuse of these medications has led to the development of antimicrobial resistance, limiting treatment options. Currently, there is growing interest in exploring the antimicrobial potential of medicinal plant extracts as an effective alternative to conventional drugs.

**Objective:** This study aimed to assess the antimicrobial capabilities of S. arborescens and S. bahorucana plants against microorganisms isolated from patients exhibiting clinical symptoms suggestive of otitis.

**Materials and Methods:** A cross-sectional, in vitro experimental study was conducted, involving the preparation of extracts from Salvia arborescens and Salvia bahorucana using various organic solvents. We used a modified agar dilution method to test the antimicrobial activity against ATCC microorganisms and clinical strains. The phytochemical composition of the plants was determined using thin-layer chromatography (TLC).

**Results:** Extracts from S. arborescens exhibited a significant presence of flavonoids and saponins. In contrast, S. bahorucana extracts displayed mild to moderate traces of the analyzed phytochemicals. Both plant extracts demonstrated antimicrobial activity capable of inhibiting the growth of strains, including S. aureus ATCC, S. pyogenes ATCC, and P. aeruginosa ATCC. The ethanolic extracts exhibited satisfactory activity against clinical isolate strains, effectively inhibiting the growth of all tested microorganisms. Notably, the ethanolic extract of S. arborescens exhibited stronger antimicrobial activity.

**Conclusions:** The results of this study provide a foundation for further research aimed at exploring the mechanisms of action and safety of medicinal plant extracts. This research may pave the way for potential clinical applications in the treatment of otitis and other infectious diseases.

## Introduction

Otitis is a pathological condition characterized by the presence of inflammation in the ear and its surrounding tissues, typically resulting from an infectious etiology [1]. Globally, the estimated prevalence of otitis is approximately 10%, encompassing an annual total of approximately 709 million occurrences. Among these instances, approximately 51% occur in children who are under the age of 5 [2]. In traditional medicine, targeted antimicrobial agents are often given to treat infections. These agents target the source of the infection, significantly improving the patient’s condition [3]. However, in recent years, the development of bacterial resistance has been linked to their improper use and overuse, which has decreased the number of therapeutic options available for treating them [4].

Currently, research into biodiversity is being conducted in an effort to find novel active ingredients in medicinal plants that have antibacterial properties. This might be a potentially useful substitute for traditional medications [5]. The World Health Organization (WHO) estimates that, due to their characteristics and the effectiveness they promise in the context of infectious disorders, 80% of the world’s population employs plants with medicinal properties for the treatment of their diseases [6]. The use of various plant-based remedies to treat infectious disorders like otitis without any supporting scientific research is a traditional practice in Latin America, which does not escape this fact [7].

In the Caribbean, there have been descriptions of using species from the *Salvia* genus for treating ear infections. This phenomenon has prompted research on plants to evaluate their antimicrobial and antifungal potential [8]. This is why the present study was created with the general objective of determining the *in vitro* antimicrobial potential of endemic plants of the Dominican Republic belonging to the genus *Salvia*, which have never been documented before, on microorganisms isolated from patients who presented a clinical picture suggesting otitis.

## Materials and methods

### Types of research

We conducted an *in vitro*, cross-sectional experimental study involving organic extracts obtained using ethanol 50%, hexane, ethyl acetate, and methanol from the endemic species *Salvia arborescens* and *Salvia bahorucana*. Subsequently, we conducted an antimicrobial susceptibility test using strains sourced from both the American Type Culture Collection (ATCC®) and clinical isolates. Additionally, we identified the phytochemical compounds present in these plants through the use of thin-layer chromatography (TLC).

### Plant collection and processing

We collected plant materials from the Valle Nuevo National Park in Constanza (*S. arborescens*) and the Sierra Bahoruco National Park (*S. bahorucana*) in the Dominican Republic. An expert in botany and taxonomy from the National Botanical Garden conducted species identification as the collection process guide. The collected material was sun-dried for 5 hours daily over a period of 7 days, subsequently ground into a powder, and stored in a cool, dry environment with proper identification for future use.

### Preparation of extracts

We obtained extracts from each plant using a ethanol 50% decoction method. Additionally, we employed the Soxhlet continuous extraction system, where 50 grams of plant material from each specimen were placed. We sequentially applied organic solvents of increasing polarity until exhaustion in the phyto-component sample (hexane, ethyl acetate, and methanol). The extractions were conducted over 6 hours daily for a duration of 3 days. We calculated the yield per gram of plant material used (see **Appendix I**). Subsequently, the extracts obtained underwent concentration via rotavaporation under reduced pressure and a controlled temperature. They were then subjected to drying in a water bath at a temperature of 40°C.

### Phytochemical analysis by Thin-Layer Chromatography (TLC)

We conducted TLC tests to identify flavonoids, saponins, coumarins, and alkaloids. The procedure involved applying 5 μL of prepared samples with a concentration of 20 mg/mL onto silica gel 60 aluminum plates. The choice of mobile phases for the tests depended on the specific secondary metabolite under investigation. We used Natural Product, Vanillin-H2SO4, and Dragendorff solutions to visualize the plates. The results were observed in a dark chamber for TLC analysis.

### Microorganisms

The microorganisms utilized in the trials included *Staphylococcus aureus* (ATCC 29212), *Streptococcus pyogenes* (ATCC 12344), *Pseudomonas aeruginosa* (ATCC 27853), *Escherichia coli* (ATCC 25922), and *Candida albicans* (ATCC 10231). Additionally, we examined four strains isolated from ear secretion cultures of patients exhibiting clinical symptoms suggestive of otitis who were admitted to a tertiary-level hospital in Santo Domingo between April and July 2022. During this period, routine antimicrobial susceptibility tests were conducted to assess the minimum inhibitory concentration (MIC). These strains were arbitrarily labeled as 001, 002, 003, and 004 for identification purposes. We also included a clinical strain of *S. aureus* (005), which had not undergone routine antimicrobial susceptibility testing by MIC and was isolated from a skin injury.

### Antimicrobial susceptibility testing

In the antimicrobial susceptibility testing, we utilized ethanol 50% extracts, which were adjusted to a concentration of 30 mg/mL. Conversely, the extracts obtained through the Soxhlet method were dissolved in a ethanol 50% solution, resulting in a concentration of 40 mg/mL.

Antimicrobial activity was assessed using the agar dilution method as described by Mitscher et al. (1972) [9]. For screening, we employed Petri plates containing Mueller-Hinton agar and the respective plant extracts at the previously mentioned concentrations. Subsequently, these plates were incubated at 37°C for 24 hours to confirm the sterility of the medium. After this initial incubation period, the ATCC and clinical microorganisms were inoculated as radial lines in duplicate and incubated at 37°C for an additional 24 hours. Gentamicin was used as a positive control at a concentration of 40 mg/mL, while ethanol at 50% served as the negative control. The results were recorded as either positive (indicating the absence of bacterial growth) or negative (indicating bacterial growth).

### Ethical considerations

We conducted all of our microbiological tests in accordance with the World Health Organization’s Guidelines for Good Practice for Pharmaceutical Microbiology Laboratories [10]. Furthermore, our protocols and evaluations adhered to the ethical standards established by the local ethics committee.

## Results

### Phytochemical analysis

In the phytochemical screening conducted via thin-layer chromatography (TLC), flavonoids were detected in small quantities within the hexane, ethyl acetate, and ethanol fractions of 80% of *S. arborescens* and in the hexane and ethyl fractions of 80% of *S. bahorucana*. Saponins were observed in the ethyl and methanol acetate fractions of both plant species. Traces of coumarins were identified in the ethanol and ethyl acetate fractions of *S. arborescens*, while *S. bahorucana* exhibited minimal traces of this phytochemical in its ethyl acetate fraction. Alkaloids were discerned in the various extracts of *S. arborescens*, except for the methanolic extract. Conversely, alkaloids were not found in any of the tested fractions of *S. bahorucana* (see **Table 1**).

**Table 1:** Phytochemical analysis results for *S. arborescens* and *S. bahorucana* using TLC.

| Phytochemicals | Mobile phase | Developing reagent | Interpretation | Extract | *S. arborescens* | *S. bahorucana* |
| --- | --- | --- | --- | --- | --- | --- |
| Flavonoids | EtOAc:Formic:HOAc:H<sub>2</sub>O (100:11:1:27) | Natural product+UV-365nm | Blue, yellow, white, fluorescent green, and purple | Hexane | ++ | ++ |
|  |  |  |  | Ethyl acetate | ++ | - |
|  |  |  |  | Methanol | - | - |
|  |  |  |  | Ethanol 80% | ++ | ++ |
| Saponins | Chloroform:MeOH:H<sub>2</sub>O (65:50:10) | Vanillin-H<sub>2</sub>SO<sub>4</sub> | Purple, red and chocolate | Hexane | - | - |
|  |  |  |  | Ethyl acetate | +++ | + |
|  |  |  |  | Methanol | +++ | + |
|  |  |  |  | Ethanol 80% | ++ | - |
| Coumarins | EtOAc (100) | NP/PEG+UV365nm | Blue and green | Hexane | + | - |
|  |  |  |  | Ethyl acetate | +++ | + |
|  |  |  |  | Methanol | - | - |
|  |  |  |  | Ethanol 80% | ++ | - |
| Alkaloids | EtOAc:MeOH:H<sub>2</sub>O (100:13.5:10) | Dragendorff+UV 254nm+UV 365 nm | Blue and yellow | Hexane | + | - |
|  |  |  |  | Ethyl acetate | + | - |
|  |  |  |  | Methanol | - | - |
|  |  |  |  | Ethanol 80% | + | - |

**Legend:** +: Slight presence; ++: Moderate presence; +++: Abundant presence; -: Absent.

### Isolation of microorganisms

A total of 39 samples were collected between April and July 2022. Twelve of them showed growth, giving a positive rate of 30.76% in ear secretion cultures from people who had symptoms suggestive of otitis. The most commonly identified microorganism was *Pseudomonas aeruginosa* (see **Figure 1**).

**Figure 1:**
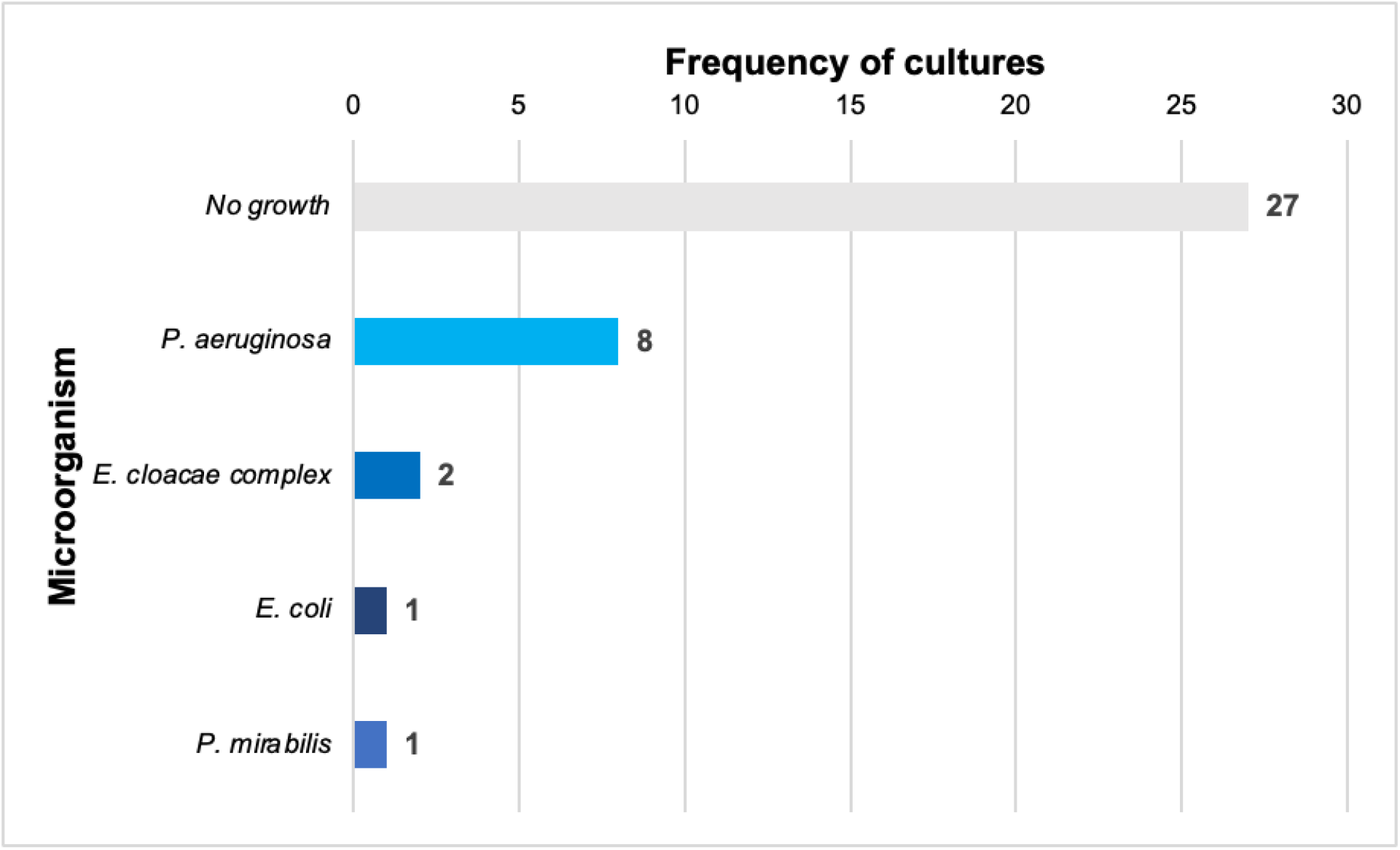
Microorganisms isolated in ear secretion cultures of patients with clinical symptoms suggestive of otitis, April–July 2022.

### Antimicrobial susceptibility tests

The standard antibiotics used, cefazolin and tigecycline, had the expected minimum inhibitory concentration in the clinical species that were tested. Resistance to ciprofloxacin was observed in one strain of *P. aeruginosa* (003) and in *S. aureus* (005) (see **Table 2**).

**Table 2:** Susceptibility test of isolated bacterial strains.

| Antimicrobial susceptibility |  |  |  |
| --- | --- | --- | --- |
| Isolated microorganism | Antimicrobial | MIC (µg/ml) | Susceptibility |
| *Pseudomonas aeruginosa* (001) | Cefazoline | ≥64 | R |
|  | Cefepime | 2 | S |
|  | Ciprofloxacin | ≤0.06 | S |
|  | Gentamicin | ≤1 | S |
|  | Tigecycline | 1 | R |
| *Pseudomonas aeruginosa* (002) | Cefazoline | 8 | R |
|  | Cefepime | 1 | S |
|  | Ciprofloxacin | ≤0.06 | S |
|  | Gentamicin | ≤1 | S |
|  | Tigecycline | ≥8 | R |
| *Pseudomonas aeruginosa* (003) | Cefazoline | ≥64 | R |
|  | Cefepime | 4 | S |
|  | Ciprofloxacin | ≥4 | R |
|  | Gentamicin | 2 | S |
|  | Tigecycline | ≥8 | R |
| *Enterobacter cloacae* complex (004) | Cefazoline | ≥64 | R |
|  | Cefepime | ≤0.12 | S |
|  | Ciprofloxacin | ≤0,06 | S |
|  | Gentamicin | ≤1 | S |
|  | Tigecycline | ≤0.5 | S |
| *Staphylococcus aureus* (005) | Gentamicin | - | S |
|  | Ciprofloxacin | - | R |

**Legend:** R: Resistant; S: Sensitivity.

### Antimicrobial activity of plants against microorganisms associated with otitis

The results of this study suggest that the species *S. arborescens* possesses significant antimicrobial properties against certain ATCC microorganisms. The ethanol 50% extract stopped the growth of *S. aureus* and *P. aeruginosa* completely. For *S*.

*pyogenes*, it inhibited growth by 80%. Ethyl acetate and methanol extracts also demonstrated a 100% inhibition in the growth of *S. pyogenes* and *S. aureus*, while the hexane extract successfully inhibited the growth of *S. aureus*. It is important to note, however, that *S. arborescens* extracts did not exhibit antimicrobial activity against *E. coli* and *C. albicans* (see **Table 3**; **Figure 2**).

**Table 3:** Antimicrobial activity of organic extracts from *Salvia arborescens* against ATCC microorganisms.

| Extracts<br>(40 mg/ml) | Microorganisms |  |  |  |  |
| --- | --- | --- | --- | --- | --- |
|  | **S.**<br>***pyogenes***<br>(ATCC<br>19615)<br>(IG%) | **E.**<br>***coli***<br>(ATCC<br>25922)<br>(IG%) | **S.**<br>***aureus***<br>(ATCC<br>25923)<br>(IG%) | **P.**<br>***aeruginosa***<br>(ATCC<br>27853)<br>(IG%) | **C.**<br>***albicans***<br>(ATCC<br>10231)<br>(IG%) |
| ***S. arborescens***<br>Hexane | 0 | 0 | 100 | 0 | 0 |
| ***S. arborescens***<br>Ethyl acetate | 100 | 0 | 100 | 0 | 0 |
| ***S. arborescens***<br>Methanol | 100 | 0 | 100 | 0 | 0 |
| ***S. arborescens***<br>Ethanol 50% | 80 | 0 | 100 | 100 | 0 |

**Legend:** IG: Inhibition growth; 0%: Total growth; 50-80%: Partial growth; 100%: No growth.

**Figure 2:**
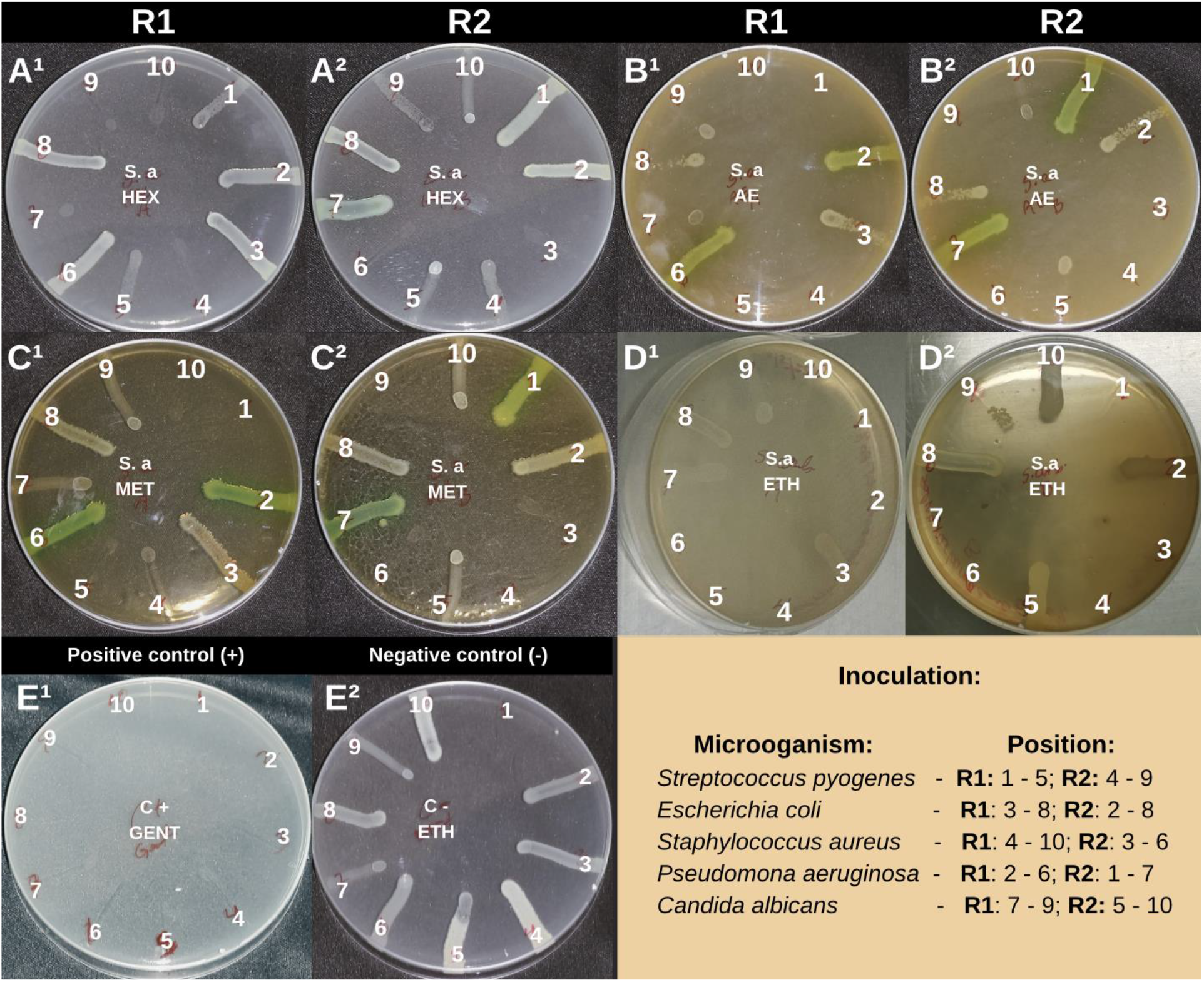
Antimicrobial activity assay of organic extracts from *Salvia arborescens* against ATCC microorganisms. **Legend:** R1: Repetition #1; R2: Repetition #2; A1-A2: Hexane (HEX); B1-B2: Ethyl acetate (AE); C1-C2: Methanol (MET); D1-D2: Ethanol 50% (ETH); E1: Positive control (GENT = Gentamicin); E2: Negative control (ETH); n= 5.

The ethanol 50% extracts of *S. bahorucana* were capable of inhibiting the growth of ATCC strains of *S. pyogenes* and *S. aureus* by 80% and 100%, respectively. Furthermore, ethyl acetate and methanol extracts achieved 100% growth inhibition against *S. pyogenes* and *S. aureus*, while the hexane extract effectively inhibited the *S. aureus* strain. No antimicrobial effects were observed against *P. aeruginosa, E. coli*, or *C. albicans*. (see **Table 4**; **Figure 3**).

**Table 4:** Antimicrobial activity of organic extracts from *Salvia bahorucana* against ATCC microorganisms.

| Extracts<br>(40 mg/ml) | Microorganisms |  |  |  |  |
| --- | --- | --- | --- | --- | --- |
|  | ***S. pyogenes***<br>(ATCC 19615)<br>(IG%) | ***E. coli***<br>(ATCC 25922)<br>(IG%) | ***S. aureus***<br>(ATCC 25923)<br>(IG%) | ***P. aeruginosa***<br>(ATCC 27853)<br>(IG%) | ***C. albicans***<br>(ATCC 10231)<br>(IG%) |
| ***S. baborucana***<br>Hexane | 0 | 0 | 100 | 0 | 0 |
| ***S. baborucana***<br>Ethyl acetate | 100 | 0 | 100 | 0 | 0 |
| ***S. baborucana***<br>Methanol | 100 | 0 | 100 | 0 | 0 |
| ***S. baborucana***<br>Ethanol 50% | 80 | 0 | 100 | 0 | 0 |

**Legend:** IG: Inhibition growth; 0%: Total growth; 50-80%: Partial growth; 100%: No growth.

**Figure 3:**
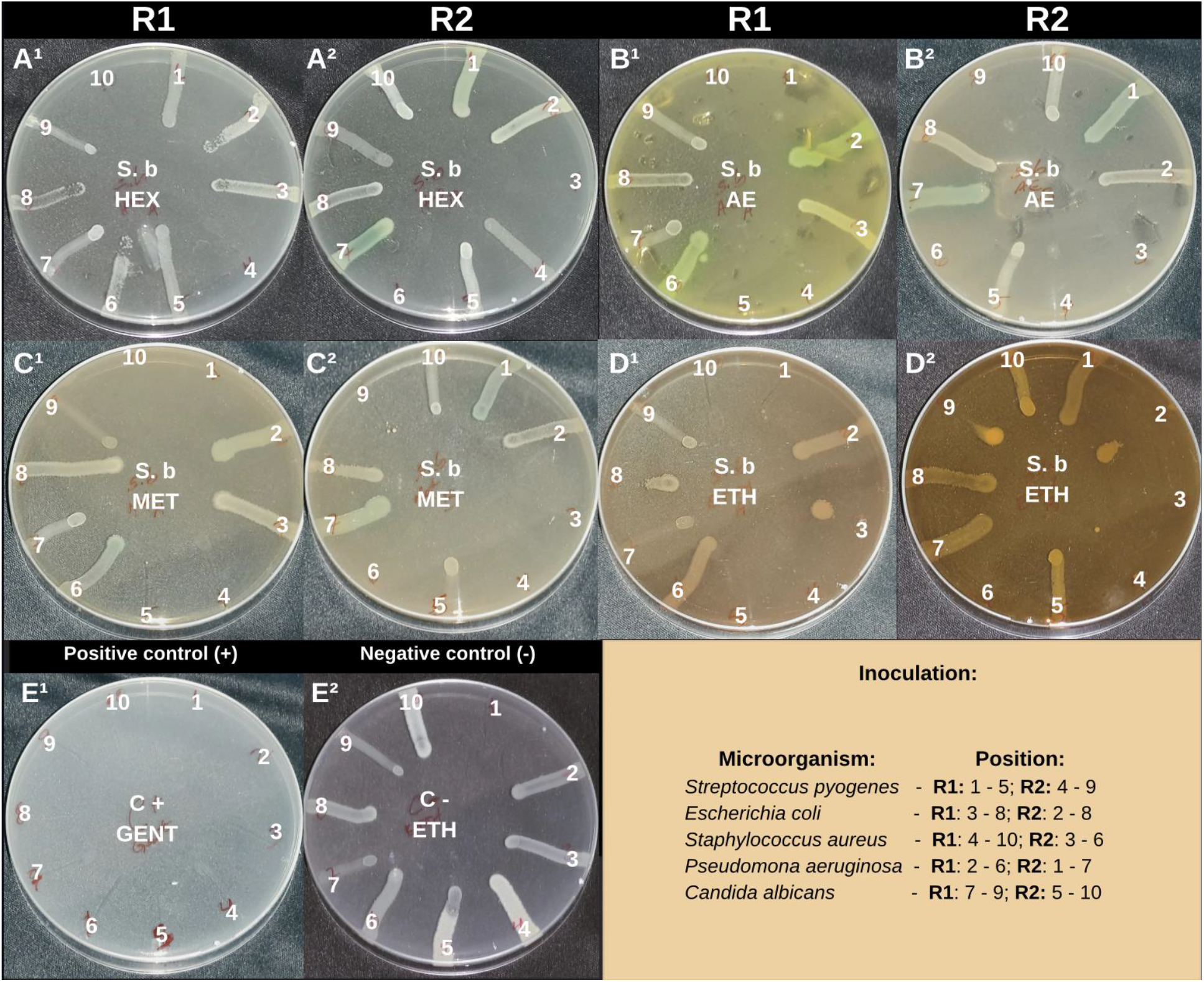
Antimicrobial activity assay of organic extracts from *Salvia bahorucana* against ATCC microorganisms. **Legend:** R1: Repetition #1; R2: Repetition #2; A1-A2: Hexane (HEX); B1-B2: Ethyl acetate (AE); C1-C2: Methanol (MET); D1-D2: Ethanol 50% (ETH); E1: Positive control (GENT = Gentamicin); E2: Negative control (ETH); n= 5.

The ethanol 50% extract from *S. arborescens* demonstrated a 100% growth inhibition in the tested clinical strains, except for the *P. aeruginosa* strain (002), which exhibited an inhibition percentage not exceeding 80% (see **Table 5**; **Figure 4**).

**Table 5:** Antimicrobial activity of organic extracts from *Salvia arborescens* against clinical otitis-associated microorganisms.

| Extract | Microorganisms |  |  |  |  |
| --- | --- | --- | --- | --- | --- |
|  | *P. aeruginosa* 001 (IG%) | *P. aeruginosa* 002 (IG%) | *P. aeruginosa* 003 (IG%) | *E. Cloacae* 004 (IG%) | *S. aureus* 005 (IG%) |
| **S. arborescens**<br>Ethanol 50% | 100 | 50 | 100 | 100 | 100 |
| **S. bahorucana**<br>Ethanol 50% | 100 | 100 | 100 | 100 | 100 |

**Legend:** IG: Inhibition growth; 0%: Total growth; 50-80%: Partial growth; 100%: No growth.

**Figure 4:**
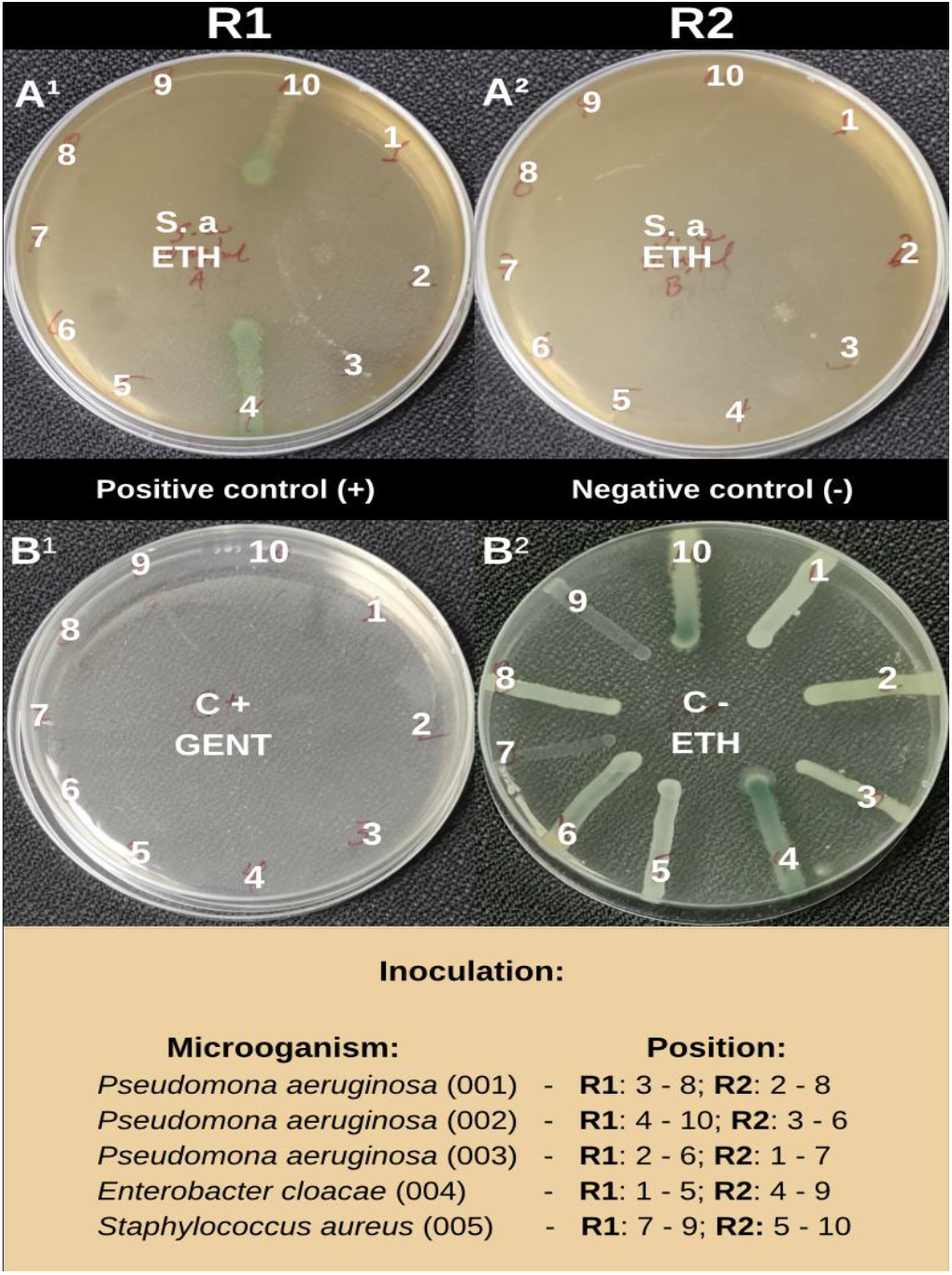
Antimicrobial activity assay of organic extracts from *Salvia arborescens* on clinical otitis-associated microorganisms. **Legend:** R1: Repetition #1; R2: Repetition #2. A1-A2: 50% Ethanol (ETH); B1: Positive control (GENT = gentamicin); B2: Negative control (ETH); n= 5.

The ethanol 50% extract of *S. bahorucana* resulted in 100% growth inhibition in all the tested strains (see **Table 6**; **Figure 5**).

**Table 6:**
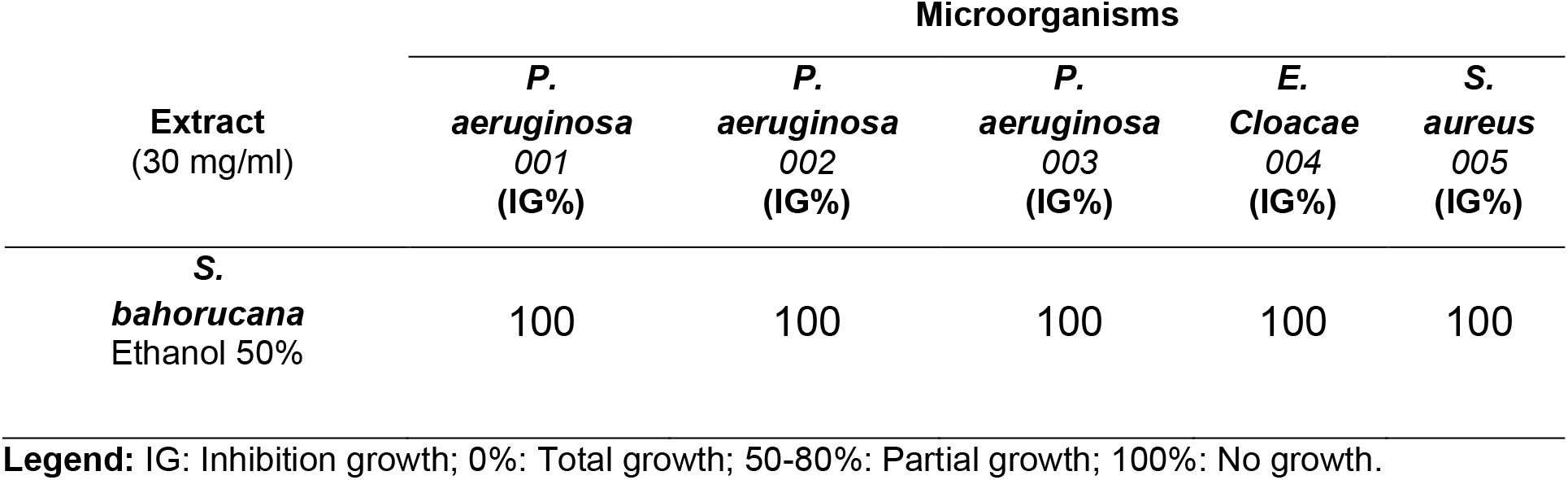
Antimicrobial activity of organic extracts from *Salvia bahorucana* against clinical otitis-associated microorganisms.

| Extract<br>(30 mg/ml) | Microorganisms |  |  |  |  |
| --- | --- | --- | --- | --- | --- |
|  | *P.*<br>*aeruginosa*<br>001<br>(IG%) | *P.*<br>*aeruginosa*<br>002<br>(IG%) | *P.*<br>*aeruginosa*<br>003<br>(IG%) | *E.*<br>*Cloacae*<br>004<br>(IG%) | *S.*<br>*aureus*<br>005<br>(IG%) |
| **S.**<br>**bahorucana**<br>Ethanol 50% | 100 | 100 | 100 | 100 | 100 |

**Legend:** IG: Inhibition growth; 0%: Total growth; 50-80%: Partial growth; 100%: No growth.

**Figure 5:**
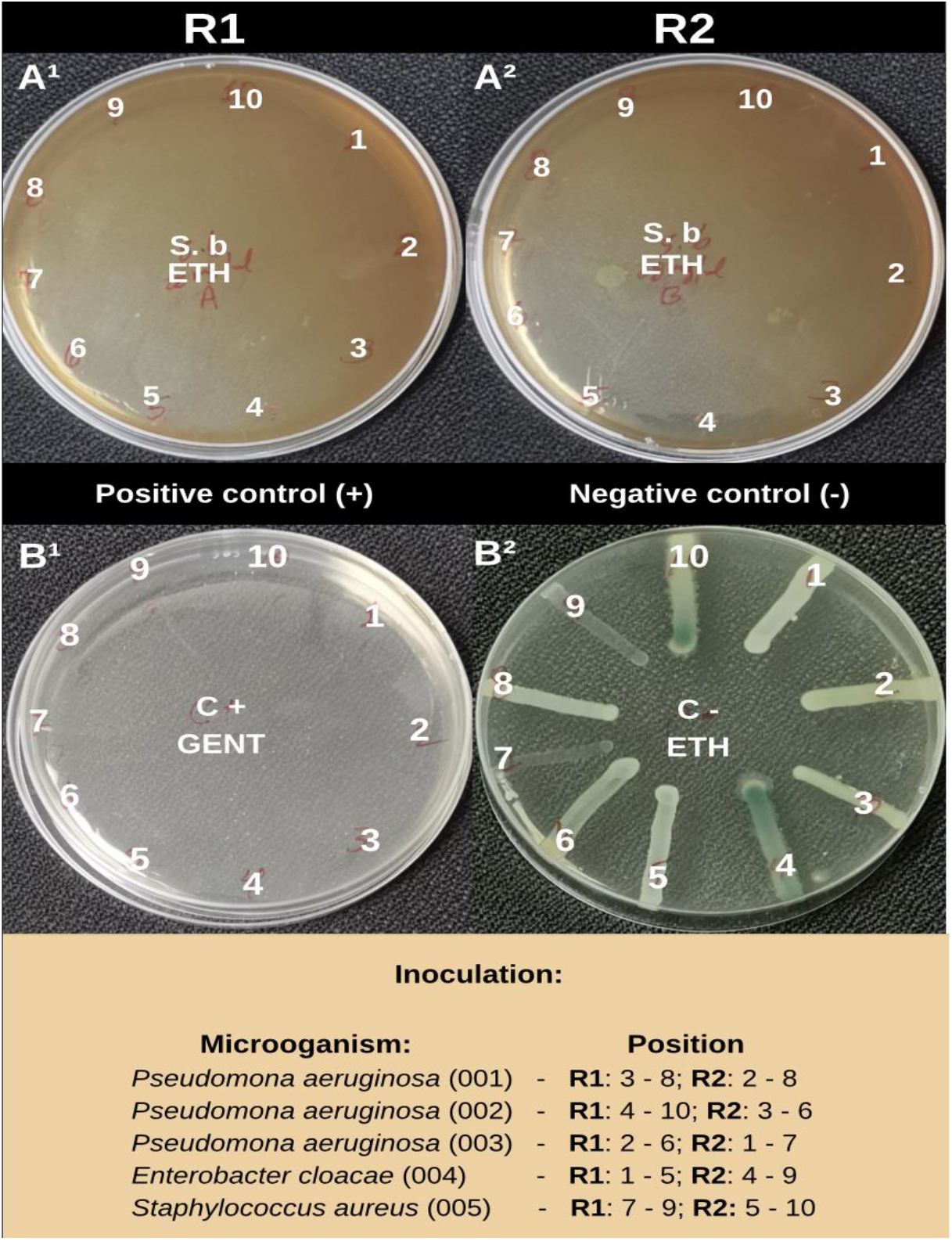
Antimicrobial activity assay of organic extracts from *Salvia bahorucana* on clinical otitis-associated microorganisms. **Legend:** R1: Repetition #1; R2: Repetition #2. A1-A2: 50% Ethanol (ETH); B1: Positive control (GENT = gentamicin); B2: Negative control (ETH); n= 5.

## Discussion

Medicinal plants have been used as a source of raw materials for making medicines since ancient times. They are still seen as an effective alternative for treating a wide range of medical conditions in many fields, including otolaryngology [7].

To our knowledge, this study is the first to demonstrate the antimicrobial activity of the endemic species *S. arborescens* and *S. bahorucana* against microorganisms associated with otic infections. Additionally, it provides valuable insights into their phytochemical composition. The phytochemical analysis unveiled a distinct phytochemical profile for each plant, despite their shared phylogenetic genus.

The analysis of S. arborescens extracts revealed the abundant presence of flavonoids and saponins, along with moderate traces of coumarins and alkaloids. Conversely, S. bahorucana extracts exhibited mild to moderate traces of the analyzed phytochemicals.

Otitis is a prevalent condition in primary care consultations, with *S. aureus* and *P. aeruginosa* being the most frequently implicated microorganisms in cases of otitis externa [3, 13, 14]. Interestingly, this study identified *P. aeruginosa* as the most commonly isolated microorganism, followed by the *E. cloacae complex*. It’s worth noting that the *E. cloacae* complex is an infrequent cause of otitis externa [13]. The isolation of the *E. cloacae* complex may be associated with various factors, including conditions like immunosuppression or extended hospitalization.

The susceptibility profile of the microorganisms to the standard antimicrobials used in our tests demonstrated minimum inhibitory concentrations within the expected parameters. However, resistance to ciprofloxacin was noted in the cases of *P. aeruginosa* (003) and *S. aureus* (005).

Despite variations in the phytochemical composition of the two plants, our tests revealed significant antimicrobial activity. This activity might be attributed to the presence, albeit in varying degrees, of various phytochemicals recognized for their antibacterial and antifungal properties. These findings are in agreement with previous studies that have highlighted the therapeutic potential of *Salvia* species [8, 11].

Our research results reveal the inhibitory activity of *S. arborescens* against both gram-positive and gram-negative microorganisms, while *S. bahorucana* primarily exhibited activity against gram-positive microorganisms. The variation in inhibitory activity between the extracts tested on the ATCC microorganisms and those on the isolated clinical microorganisms may be attributed to differences in the extract preparation methods.

## Conclusion

The findings of this research provide a solid foundation for future investigations aimed at delving into the mechanisms of action and safety of medicinal plant extracts. Such studies could pave the way for potential clinical applications in the treatment of otitis and other infectious diseases, as well as offer new avenues for combating antimicrobial resistance.

The phytochemical analysis revealed that S. arborescens extracts contained an abundance of flavonoids and saponins, along with moderate traces of coumarins and alkaloids. Conversely, S. bahorucana extracts exhibited mild to moderate traces of the analyzed phytochemicals. Throughout the study period, the most frequently isolated microorganisms were P. aeruginosa and the E. cloacae complex.

The extracts from both plants displayed notable antimicrobial activity, effectively inhibiting the growth of strains from quality control collections and clinical isolates. Notably, the ethanolic extracts exhibited satisfactory performance against clinical strains, successfully inhibiting the growth of all microorganisms tested. Among the extracts, the ethanolic extract from S. arborescens demonstrated the most potent antimicrobial activity.

However, it is essential to highlight that further investigations are warranted to gain a deeper understanding of the mechanisms of action and safety associated with these extracts before contemplating their clinical applications.

## Appendix

**Appendix 1:**
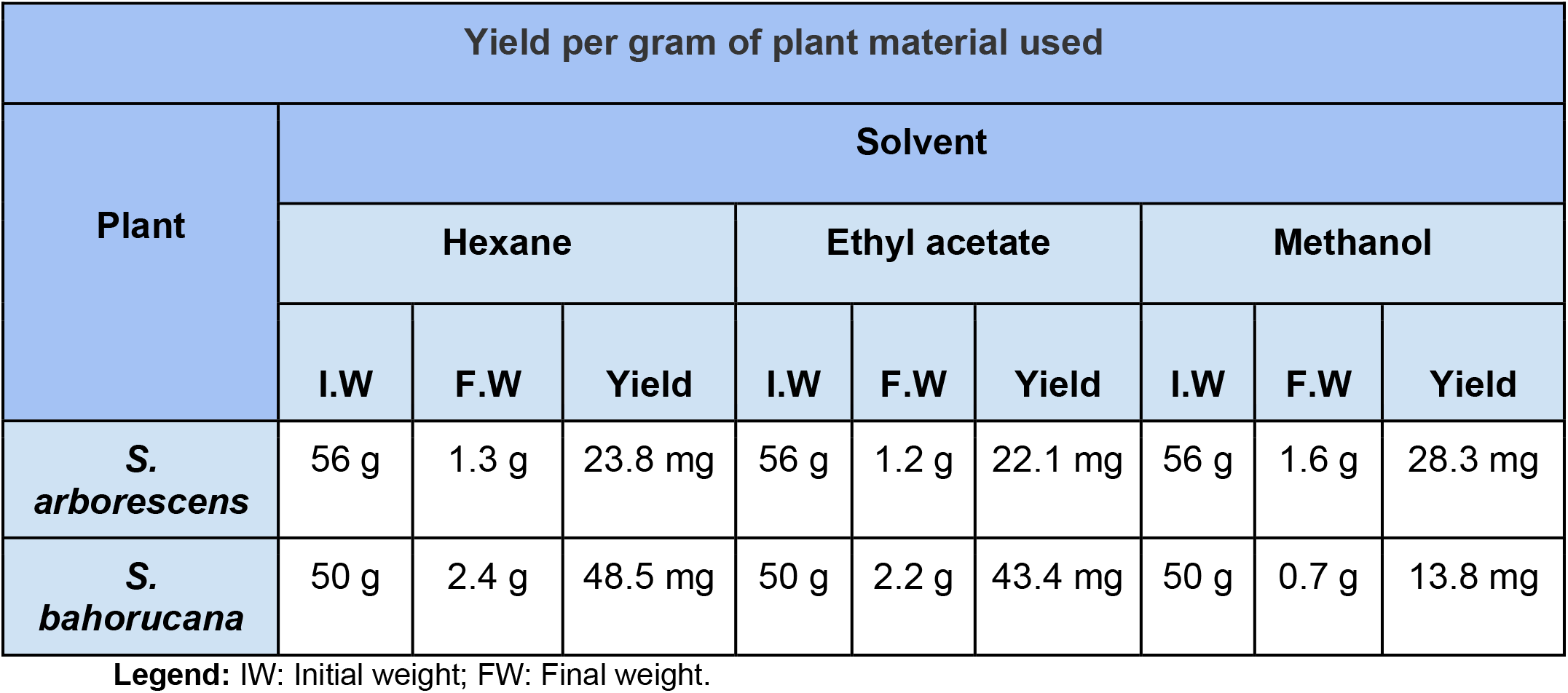
Yield per gram of plant material used.

| Yield per gram of plant material used |  |  |  |  |  |  |  |  |  |
| --- | --- | --- | --- | --- | --- | --- | --- | --- | --- |
| Plant | Solvent |  |  |  |  |  |  |  |  |
|  | Hexane |  |  | Ethyl acetate |  |  | Methanol |  |  |
|  | I.W | F.W | Yield | I.W | F.W | Yield | I.W | F.W | Yield |
| **S.**<br>***arborescens*** | 56 g | 1.3 g | 23.8 mg | 56 g | 1.2 g | 22.1 mg | 56 g | 1.6 g | 28.3 mg |
| **S.**<br>***bahorucana*** | 50 g | 2.4 g | 48.5 mg | 50 g | 2.2 g | 43.4 mg | 50 g | 0.7 g | 13.8 mg |

**Legend:** IW: Initial weight; FW: Final weight.

